# Chromatin Capture Upsampling Toolbox - CCUT: A Versatile and unified Framework to Train Your Chromatin Capture Deep Learning Models

**DOI:** 10.1101/2024.05.29.596528

**Authors:** Stanislav Sys, Alejandro Ceron-Noriega, Anne Kerber, Stephan Weißbach, Susann Schweiger, Michael Wand, Karin Everschor-Sitte, Susanne Gerber

## Abstract

Chromatin Capture Experiments such as Hi-C and Micro-C have become popular methods for genome architecture exploration. Recently, also a protocol for long read sequencing, Pore-C, was introduced, allowing the characterization of three-dimensional chromatin structures using Oxford Nanopore Sequencing Technology. Here, we present a framework that focuses on the efficient reconstruction of low-resolution Pore-C data but can also process all other 3C data, such as Hi-C and Micro-C matrices, using models that can be trained on a consumer GPU. Furthermore, we integrate building blocks of popular super-resolution methods such as SWIN-Transformer or residual-in-residual-blocks to modify or build customized networks on the fly. Pre-built models were trained and evaluated on multiple publicly available gold-standard Micro-C and Pore-C datasets, allowing for fine-scale structure prediction. Our work aims to overcome the drawback of high sequencing costs to construct high resolution contact matrices, as well as the problem of mapping low-coverage libraries to high-resolution structures in the genome. Although there have been major breakthroughs regarding NGS-based methods for the reconstruction of high-resolution chromatin interaction matrices from low-resolution data, for data obtained by long-read sequencing, there is currently no solution to reconstruct missing and sparse information and to improve the quality.

**Availability:** The tool is available at (https://github.com/stasys-hub/CCUT)

## Introduction

Chromatin Conformation Capture techniques (3C) such as Hi-C and Micro-C have been extensively utilized to investigate higher-order interactions in genome architecture [1], [2]. Such methods measure the interaction frequencies between different loci in the genome and can be used to locate specific structures such as A/B compartments, chromatin loops, or topologically associated Domains (TADs) in the genome. These secondary DNA structure features significantly influence gene expression via its spatial configuration within the nucleus [3], [4]. More recent methods such as Micro-C [5] allow for a more fine-grained picture of the spatial organization of chromatin in the nucleus. The reconstruction of the three-dimensional arrangement of these components is essential for deciphering the relationships between chromatin organization and the genetic function [1], [2], [4]–[7]. These techniques also play a key role in the detection of structural variants (SVs) that disrupt chromatin’s three-dimensional structure, thereby affecting gene expression. Such alterations can subsequently lead to diseases, including cancer, through mechanisms such as the disruption of tumor suppressor genes, the duplication of proto-oncogenes, or the formation of oncogenic fusion genes. [3], [6]–[8]. Despite all the advances achieved using these techniques, 3C-based methods still face significant obstacles in the characterization of higher-order chromatin structures. Underlying these problems are the typical limitations of short-read sequencing, which result in uneven coverage of the genome in regions with different GC content, reduced detection of large-scale DNA rearrangements and challenges in mapping sequences within repetitive DNA regions [9]–[13]. These challenges are discussed in detail by excellent reviews from by Li and Durbin as well as Zhang et al [14], [15]. This gap has recently been closed by the Pore-C protocol from Oxford Nanopore [9], [12], [16], [17]. Pore-C provides an additional layer of information compared to the classical 3C methods by providing access to global high-order multiway contacts and capturing crucial methylation information at the same time [18]. These advancements are fundamental, as gene regulation often entails intricate chromatin interactions across multiple loci, linking enhancers and promoters [4], [19]. Furthermore, the structural arrangement of chromatin exhibits cellular and contextual specificity, thereby introducing an additional layer of complexity in leveraging publicly accessible datasets [5], [20]. However, the performance of the algorithms used to map such data to genome organization and the quality of their predictions, correlate highly with the data quality, especially the amount of reads produced, which directly translates to the maximum possible resolution for pairwise interaction matrices. Generating matrices of high quality with a resolution ≤ 10kb presents challenges in both production feasibility and effectiveness, leading to increased costs until satisfactory results are obtained. This is primarily due to the quadratic increase in the required number of reads necessary to achieve only a linear enhancement in resolution. Publicly available data can be used for reconstruction and qualitative improvement under certain conditions but their availability is sparse. For example, to obtain the entire three-dimensional structure of a complete human genome at high resolution using Pore-C, typically at least four PromethION flow cells must be used per genome [21]. The need to generate such an amount of data increases both the cost and the labour intensity, highlighting the need for methods that can overcome these challenges in both technologies. To address this shortcoming for NGS-based 3C data, multiple tools have been developed [22]–[28]. The majority of them are based on deep learning approaches and extrapolate popular techniques from computer vision for image reconstruction as adversarial training and convolutional neural networks (CNNs). However, these models possess limitations that require detailed tuning or retraining because they depend on specific training data, for example working only with normalized data or assuming precentile cut offs to deal with outliers. While especially generative adversarial neural network (GAN) based approaches provide excellent reconstruction results, the training process can be unstable and computationally demanding [29], requiring month-long computing time on expensive professional grade hardware, which most biological labs lack. Furthermore, all these methods diverge significantly in their assumptions about the data and its transformation. Due to the absence of standardized protocols and benchmarks for such tools, methodological comparisons become very challenging, leading to results that can be irreproducible when applied to new but similar datasets, especially when solely relying on common metrics from computer vision as shown by Yamashita et al. [30], [31]. For Pore-C (long-read 3C) data, there are currently no established methods for improving the data, which is a major obstacle to effective chromatin reconstructions using nanopore sequencing. Here, we introduce the Chromatin Capture Upsampling Toolbox (CCUT), showcasing its deployment of cost-efficient restoration models pre-trained on gold-standard, publicly accessible datasets (see Fig. 1). Moreover, CCUT does not only contain pre-built models but also encourages to modify and build models in a unified manner. To this end, we provide unified base classes and deep learning building blocks as well as data loaders to increase reproducibility and make custom models more accessible. Furthermore, we include utilities to visualize and transform data. The toolbox was designed to be extendable, empowering users to customize the functionality to their needs. To demonstrate the capabilities of CCUT we built and trained multiple models on Deshpande et al.’s gold-standard Pore-C datasets [21], as well as on classical 3C datasets. Moreover, we evaluate the performance of restoration models with respect to the applied data augmentations, optimal kernel sizes, and loss functions. We propose a novel Fast-Fourier-Transform inspired loss function for our models, which considerably enhances the visibility of existing chromatin structures. To simulate low-resolution data, we downsampled the high-quality datasets on the read level by the factors of 16x and 100×. Since TADs represent the most common form of spatial interaction in the genome and have an average length of 1 mio bp, we aimed for a target resolution of 200×200 pixels, where every pixel represents a 10kb bin, leading to the coverage of a 2 mio bp window. Hereby, our models are able to match or even improve on accuracy compared to current models as HiCARN or ScHiCEDRN [23], [28] and impute fine-grained genomic structures. Additionally, we conduct a case study on both gold standard Pore-C data and Micro-C data; the former presents unique challenges and serves as a novel element in our study due to its distinct characteristics.

**Figure 1:**
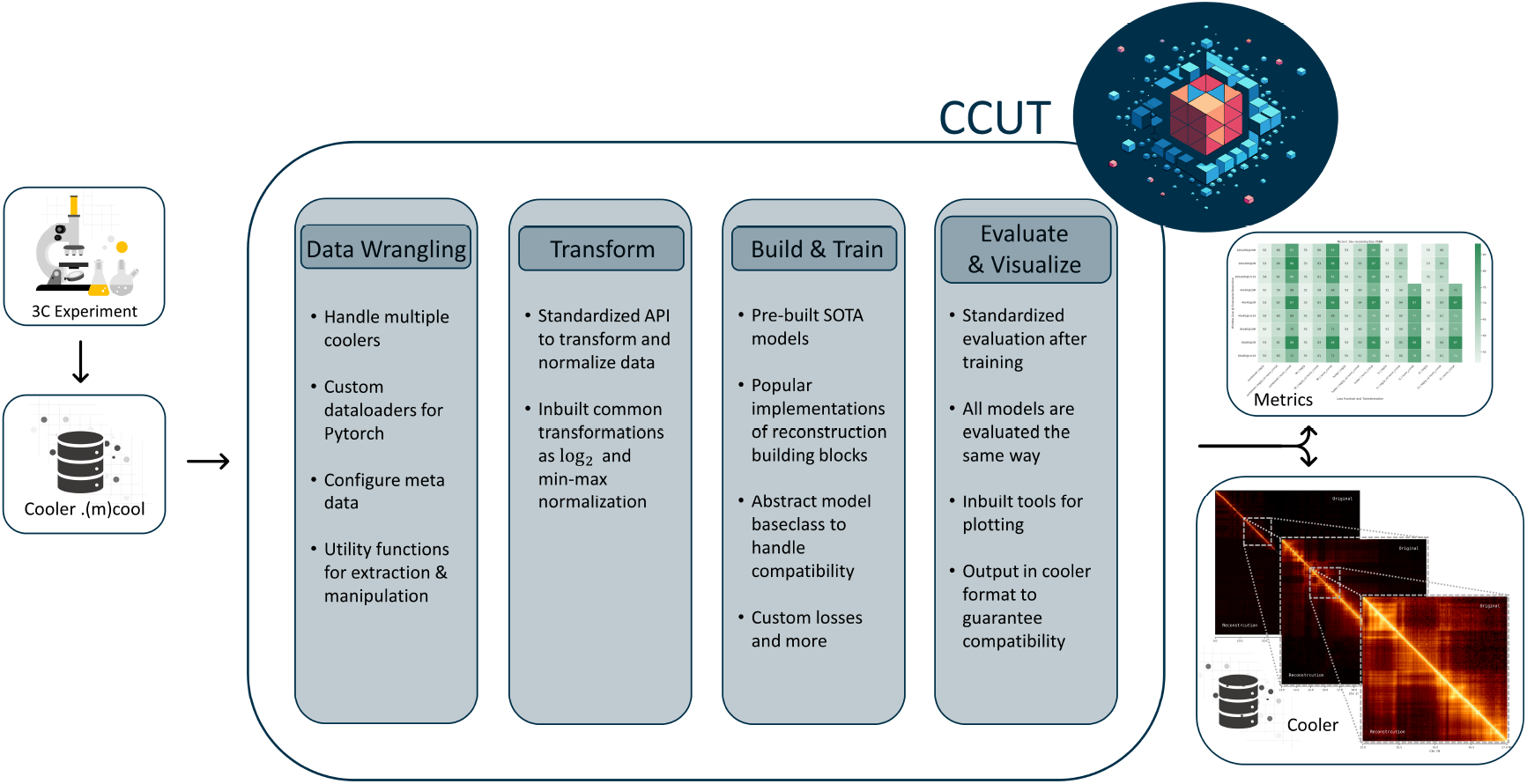
Overview of CCUT. A versatile deep learning toolbox built upon Pytorch to reconstruct and enhance 3C-contact matrices. Key components include: (1) Data Wrangling, facilitates the handling of multiple Cooler file formats and includes custom PyTorch dataloaders as well as utility functions for data extraction and manipulation. (2) Transform, features a unified API that implements common transformations such as log_2_ and min-max normalization for data preprocessing. (3) Build & Train, includes pre-built state-of-the-art (SOTA) model architectures, reconstruction algorithm implementations, an abstract model base class for compatibility, and custom loss functions. (4) Evaluate & Visualize, offers standardized post-training evaluation, plotting tools, and output in Cooler format to ensure compatibility across analyses. Metrics are systematically collected for model assessment and comparison.

## Results

### Impact of Loss Functions and Data Augmentation on raw data

The impact of data transformations and model parameters on the quality of pairwise correlation matrix reconstruction lacks consensus within the scientific community. To address this, we evaluated the RRDB-UNet model on Pore-C and Micro-C datasets to measure the impact of these factors on different scales with respect to Peak Signal-to-Noise Ratio (PSNR) (see Figure 2), Structural Similarity Index Measure (SSIM), Mean Squared Error (MSE), and Mean Absolute Error (MAE). We chose to primarily report PSNR as the performance metric for our image reconstruction model, given its ease of interpretation and widespread usage across domains. PSNR effectively measures the quality of reconstructed images relative to their originals by quantifying the ratio between signal power and corrupting noise, making it well-suited for performance evaluation.

**Figure 2:**
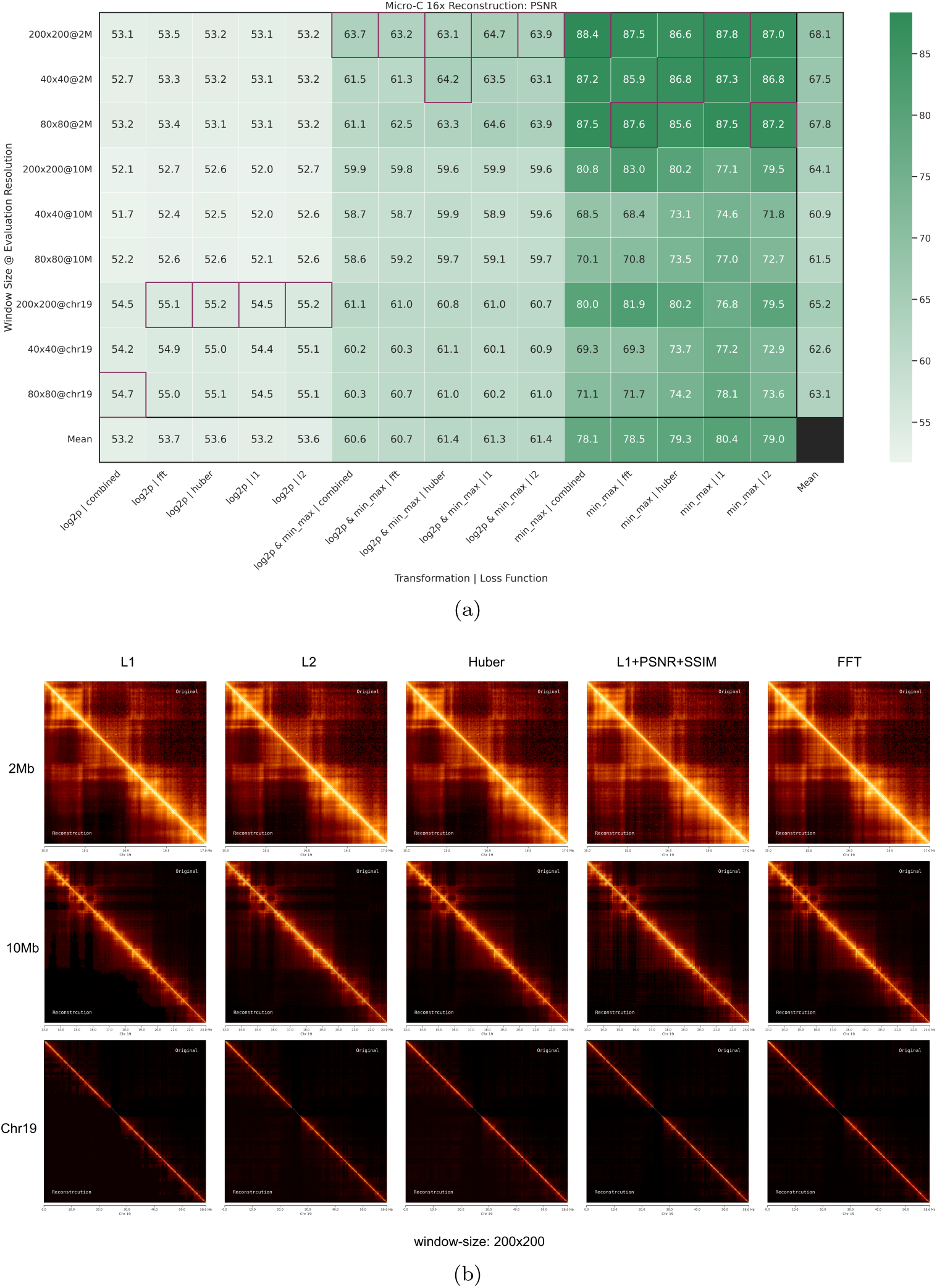
a. Heatmap representation of PSNR values across various loss functions and data transformations. Each cell corresponds to the average PSNR obtained from a model with a particular combination of loss function and transformation, across different genomic window sizes: 40×40, 80×80, and 200×200. The 200×200 window size achieves the highest scores, particularly with the combination of min-max normalization and the L1+PSNR+SSIM loss function. b. Visual comparison of log_2_ transformed original and reconstructed Micro-C contact maps with respect to different loss functions for a 2 Mb region, a 10 Mb region, and chromosome 19 (Chr19). The L1+PSNR+SSIM and FFT loss functions demonstrate a higher visual correspondence to the original images, with FFT showing the least amount of artifacts. The L1 loss notably introduces significant visual distortions, particularly in the chromosome-wide reconstruction.

Models reconstructed 2 Mb and 10 Mb regions on chromosome 19, including the chromosome itself, utilizing different patch sizes (40×40, 80×80, and 200×200). The resolution for Micro-C was set at 10 Kb and for Pore-C at 50 Kb. Models trained on 200×200 patches exhibited superior performance compared to those trained on 40×40 and 80×80 patches, displaying enhanced generalizability across various genomic scales (see Figure 2a). Larger patch sizes generally yielded higher PSNR values. However, while the 200×200 models displayed reduced reconstruction quality with increasing genomic range, they still outperformed smaller models, especially at the whole chromosome level. Smaller models performed comparably well at the smallest genomic evaluation window of 2 Mb but exhibited more rapid quality deterioration with increasing genomic intervals (see Figure 2a). This phenomenon aligns with the anticipated inverse correlation between matrix similarity and error accumulation with increased size.

#### Loss Functions

The Fast Fourier Transformation (FFT) inspired loss exhibited enhanced performance, notably in models trained on 200×200 patch sizes for reconstructing whole chromosomes. This effect was most noticeable in minmax normalized data. While FFT loss performed best in these settings, the Huber and L1 losses (see Eq. 1, 3) showed excellent generalizability across all scales evaluated. Loss functions primarily affect PSNR quality concerning min-max normalized data and tend to produce more consistent results with log-transformed data.

#### Data Transformations

The highest PSNR values are obtained with exclusive min-max normalization, regardless of the loss function employed. Models learning the min-max normalized logarithmic data also yield higher scores compared to those using solely the logarithmic transformation. Nevertheless, comparing PSNR values between differently augmented data is not feasible. One can observe that data that has been logarithmically transformed tends to have less variance when reconstructed. Model performance is nearly identical when looking at the genomic intervals of 2 and 10 Mb for the logarithmic scale. Min-max normalized raw counts display a much higher variance of reconstruction quality with respect to PSNR.

### Results of Loss Function Impact on Visual Reconstruction

In addition to classical vision metrics, we assessed reconstruction quality based on human visual perception, still considered the gold standard in computer vision ([30]). Reconstructions were evaluated with respect to their loss functions, specifically L1, L2, Huber, a synergistic combination of L1 with PSNR and SSIM (L1+PSNR+SSIM), and a Fourier Transform-based approach (FFT). These comparisons were conducted across three distinct genomic scales: a 2 Mb region, a 10 Mb region, and the entirety of chromosome 19 (chr19), all utilizing a patch size of 200×200 and all data log_2_-transformed as typical in chromatin capture analysis.

Across the different loss functions utilized, FFT-based loss function reconstructions consistently exhibit a high fidelity to the original images, with this effect being particularly pronounced in the bigger genomic intervals of 10 Mb and chr19. The reconstruction accuracy decreases as the genomic scale expands, with the 2 Mb reconstructions being markedly clearer and more artifact-free compared to the larger scales. In many cases such artifacts can only be observed with the human eye because they are results of the learning objective and can’t be measured by common distance metrics as MSE or MAE.

Notably, the L1 and L2 loss functions tend to introduce significant visual artifacts, which are especially discernible at the chromosome-wide scale where the L1 loss leads to the most pronounced artifact introduction, manifesting as darkened areas and vertical striping which are absent in the original counterpart.

Visually, the L1+PSNR+SSIM combination appears to offer an optimal balance between reconstruction precision and visual coherence across the varying scales. Upon close inspection, its performance is closely matched, if not indiscernible, from the FFT-based approach, especially when one examines the details at higher magnifications. This comparative analysis underscores the complexity of Hi-C data reconstruction, emphasizing the pivotal role of loss function selection in artifact mitigation and overall visual fidelity of reconstructed images.

#### Challenges in Chromosome-Wide Reconstruction

Our observations highlight a distinct contrast in the reconstruction quality between localized genomic regions and chromosome-wide reconstructions. Reconstructions exhibit complex interactions between the genomic scale and the loss function, particularly notable in whole chromosome reconstructions where incorrect loss function selection can result in significant visual artifacts, exemplified by the L1-Loss in Figure 2b.

### Evaluation of Pore-C & Micro-C Reconstruction

To evaluate a more realistic scenario, we calculated the insulation profiles of the reconstructed, downsampled and original matrices [32], which serve as the foundation for most TAD callers. We worked with raw count data, which was capped at the 99.9*th* percentile (317 for Micro-C, 73 for Pore-C) and normalized between 0-1 as described in [23], [25]–[28], [33]. Pore-C Data was handled at 50Kb resolution and Micro-C at 10Kb. We trained our custom UNet model for 50 epochs with early stopping if the MSE did not decrease for more than 5 epochs. Our target evaluation metric for the most performant model was SSIM, which peaked at 0.8834 for Pore-C and 0.9604 for Micro-C on average for the evaluation dataset containing chromosomes 19-22 reaching state-of-the-art reconstruction quality when compared to the newest models of ScHiCEDRN and HiCARN [23], [28]. In contrast to all previously mentioned models, which rely on the same downsampling procedure, only ScHiCEDRN, which focuses on single-cell Hi-C data, and our framework provide the compatibility to work with Cooler-files [34](see Fig. 1). Furthermore, our framework provides the opportunity to create new cooler files given non-balanced data. By leveraging this type of pre-processing, we were able to improve the contact map quality for Pore-C as well as Micro-C data on multiple scales (5Mbp, 10Mbp, 20Mbp) as depicted in Figures 3a & 3b, while also demonstrating the adaptability of our model. Despite the clear visual improvement PSNR, MSE, and SSIM improve drastically even on the smallest 5Mb scale where PSNR improves from 67.682 to 76.683, the MSE decreases from 0.011 to 0.001 and SSIM increases from 0.557 to 0.907. This holds true for the 10Mb and 20Mb scales as well as for Micro-C data, where the improvement is even more striking given that the Pore-C data was downsampled by a factor of 4x and Micro-C by a factor of 16x.

**Figure 3:**
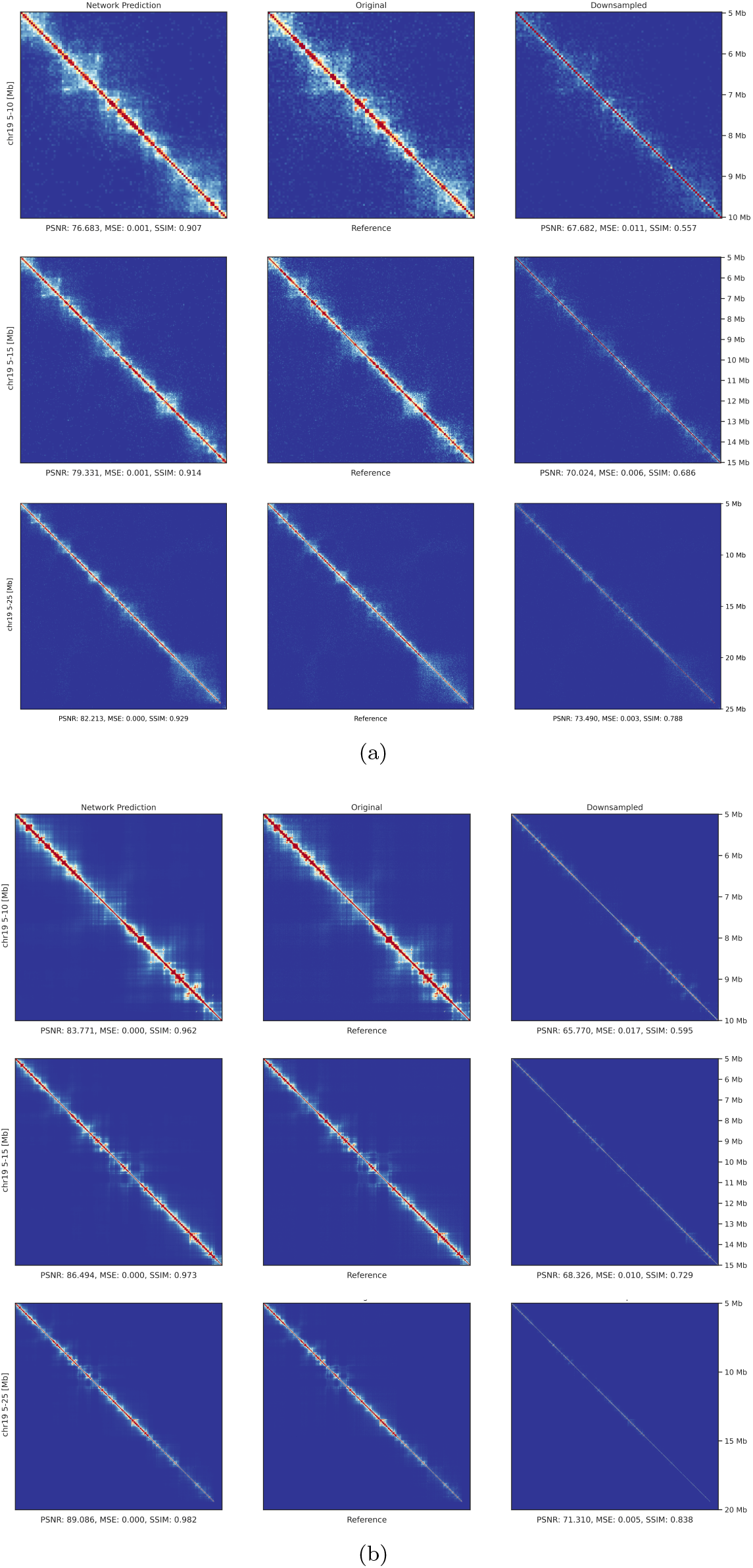
Visual comparison of chromatin interaction maps at multiple genomic scales. This figure illustrates the improvements in spatial resolution and clarity achieved through our models, highlighting both the raw and reconstructed contact maps for Pore-C (a) and Micro-C (b) datasets. Each panel per dataset displays results at 5 Mb, 10 Mb, and 20 Mb genomic intervals, showcasing the models’ effectiveness across varying levels of complexity and resolution.

#### Biological Validation

While traditional computer vision metrics such as PSNR and SSIM can give a good understanding of the quality of a visual reconstruction, they are no substitution for biological validation. To assess the similarity of 3C patterns, we first calculated Pearson’s correlation *r* across the diagonals of the matrices as described in [22] and subsequently the insulation profile as described in [32]. This profile is typically computed as a multiple of the original 3C data resolution, where we chose the factors of 3 and 5 as typical in such studies. The most common and important genomic interactions are in the form of TADs and do typically not exceed the range of 1Mb and extremely rarely 2Mb [35]–[37]; therefore, a high Pearson correlation in this interval is crucial for correct biological classification. Our Pore-C and Micro-C Reconstructions exhibit a far higher Pearson correlation in this genomic area compared to their respective downsampled counterparts (see Fig. 4). This holds true for Pore-C up to a genomic distance of ∼6Mb and for Micro-C even up to ∼36Mb, assuring that interactions below these thresholds exhibit a character more similar to high-quality data.

**Figure 4:**
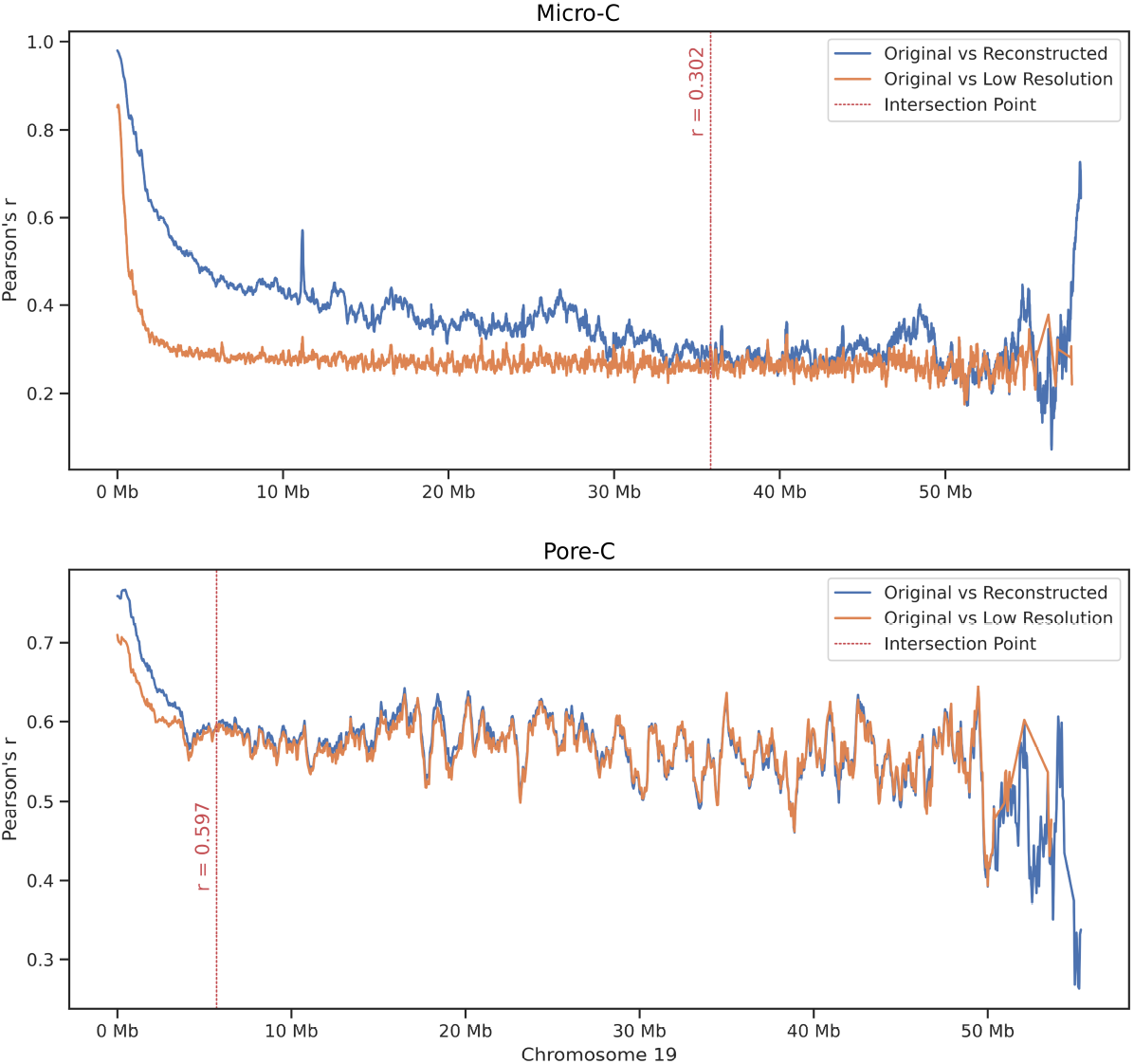
Correlation analysis of original vs. reconstructed chromatin maps. This figure presents Pearson correlation coefficient *r* calculated across the diagonals of the respective interaction matrices, comparing performance between original high-resolution data and CCUT reconstructions. The results highlight the accuracy of our models in maintaining genomic interaction patterns, with correlations illustrated at scales up to 6 Mb for Pore-C and 36 Mb for Micro-C.

To provide a measure of how accurately the biological validity of the reconstructions is captured, we calculated the MAE, MSE, RMSE, Pearson’s r, and Spearman’s *ρ* for the insulation profiles with respect to the high-quality data, as compared to our reconstructions and the downsampled data (see Fig. 5). We observe that the reconstructed insulation profiles for Hi-C and Micro-C are visually nearly indistinguishable from the original data. This observation is further supported by the standard distance metrics (MAE, MSE, and RMSE), which indicate a greater similarity and low error rates between the reconstructed profiles and the original data than between the downsampled data and the original. The Pearson correlation is higher for Micro-C when comparing downsampled to original data, whereas Spearman’s correlation is higher between the reconstruction and the original data. We observe the same trend for the Pore-C data, while only Spearman’s *ρ* is higher for 150kb profile, albeit all other metrics being in favour of the reconstructed profiles.

**Figure 5:**
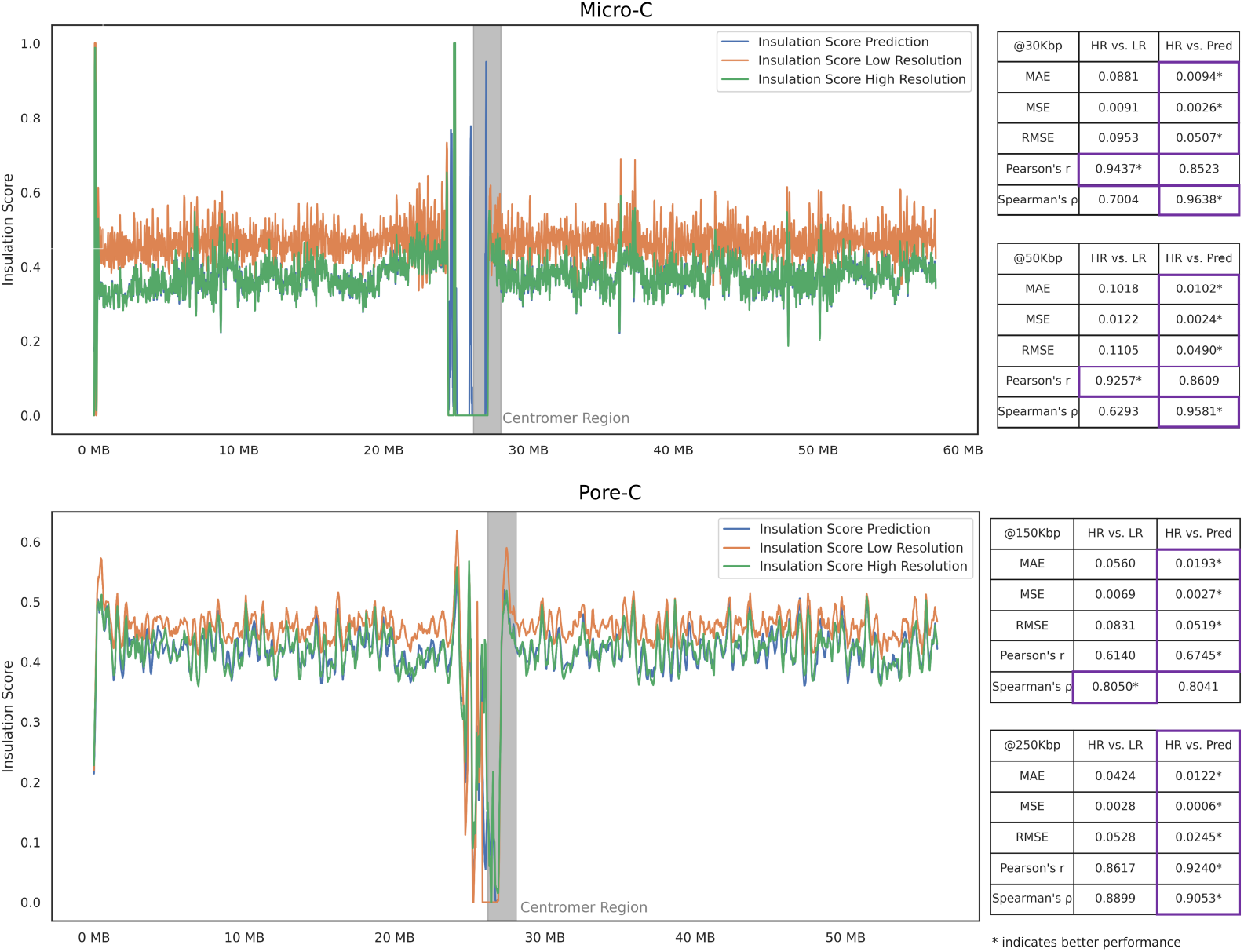
Biological validation of reconstructed interaction profiles. Displayed are insulation profiles calculated for both original and reconstructed data, alongside comparisons to downsampled datasets. The profiles demonstrate how closely our reconstructions mimic the original data, with metrics like MAE, MSE, and Pearson’s r emphasizing the high fidelity of our approach across various chromosomal distances.

## Discussion

With the fast-paced developments in deep learning and the steady advancements in computer hardware over the last few years, computational methods to assist experiments have become increasingly popular. While deep learning models offer substantial improvements in denoising and reconstructing chromatin capture data [23], [25]–[28], [33], benchmarks and best practices are yet to be established. This work tries to addresses this by providing a computational framework that enables researchers to integrate their models, use pre-trained models, or build new models using pre-built building blocks for rapid development. By providing a uniform interface for data augmentation and model training, we facilitate reproducibility and comparability between different models as shown in the application of our framework to Pore-C data, which is inherently more complex and noisy than other 3C methods (see Fig. 3a,4,5).

A core component of our study addresses the currently lacking standards in the evaluation of deep learning models for chromatin conformation capture data reconstruction. Most models, such as EnHiC, DeepHiC, or HiCARN, propose specialized model architectures with notable reconstruction results but have different assumptions for data pre-processing and transformations as percentile cut-offs or raw-count normalization. For effective benchmarking, these models must be modified/adapted to align with the criteria of the reference model used in their respective publications, despite not being originally designed/intended for this purpose [22], [23], [26]. CCUT tackles this problem by unifying this process and incorporating models like HiCARN and the respective building blocks into its model zoo, making it more straightforward to compare such parameters.

Furthermore, the efficacy of different loss functions in the reconstruction of chromatin capture data plays a critical role in model performance and influences performance in terms of visual evaluation and classical vision metrics significantly (see Fig. 2), especially when taking different scales into account rather than calculating an average metric per patch as most studies currently do [23], [25], [26], [28]. Moreover, we could show in our Benchmark (see Figure 2) that it is not trivial to compare metrics for computer vision on data that has been transformed by different methods. Fig. 2 demonstrates the impact of different loss functions, transformations, and training with different patch sizes. While patch size and loss functions have the most impact on reconstructions at bigger scales as whole chromosomes, normalization and data transformation play a key component in the quality of reconstruction. Larger window sizes can capture more complex relationships but are more expensive due to the quadratic increasing computational cost. Especially minmax normalization should be considered a cornerstone for every 3C enhancement model since it stabilizes training and boosts performance significantly. Furthermore, when chromatin capture data undergoes a logarithmic transformation, it is compressed into a much smaller interval, exhibiting significantly reduced variance compared to other normalization techniques. While this is beneficial for visual inspection, it obscures the distribution of the original count data, potentially introducing biases into downstream evaluations.

### Application to Pore-C Data

This study is the first to apply such a model to Pore-C data enhancement. While Pore-C comes with multiple advantages over existing 3C techniques, such as multi-concatemers or the ability to capture epigenetic information in parallel, its inherently noisy and complex data structure poses its own challenges for matrix reconstruction [17], [21], [38]–[40]. To mimic realistic circumstances, we simulated the performance of a less equipped ONT PromethION run, using only one flow cell instead of the typical four, by downsampling the original high-resolution Pore-C data at the read level by a factor of four. Our results show a clear visual improvement, which is underlined by the most important computer vision metrics such as PSNR and SSIM (see Fig. 3a) and clear statistical correlation between conformations (see Fig. 4). Albeit excellent statistical reconstruction, it is essential to evaluate such models based on metrics which are tailored for biological interpretation such as insulation profiles [32]. These profiles are calculated by sliding a rectangular window along the matrix to sum up contacts within a given region surrounding each locus, easily allowing inference of TADs by contact cutoffs [32]. The benefit of working with profiles rather than with TAD callers directly is that they form the basis for most TAD callers, which are a statistical construct, rather than a structural feature of the genome since their detection is hyperparameter and assumption-dependent and therefore will vary depending on the tool used [41]–[43]. Here, we could show that our tool can reconstruct matrices that produce an insulation profile more similar to the original high-quality data while keeping significantly smaller error rates (see Fig. 5). Overall, our model consistently reconstructs the original insulation profile. This setup tests the robustness of our model and its applicability to real-world scenarios where access to high-end sequencing facilities may be restricted and shows how such models can potentially be used to solve the challenge of Pore-C’s low sequencing yield compared to Illumina without having to rely on a fully loaded PromethION, by possibly reducing sequencing costs.

## Conclusion

We demonstrated that CCUT, a versatile framework designed to enhance the resolution and quality of 3C-based chromatin capture data through state-of-the-art deep learning methodologies, can help standardize training and evaluation for deep restoration models for chromatin capture data, while at the same time offering innovative new methods such as the training of models with our custom FFT loss, which performs on par with the perceptual loss. Our results highlight CCUT’s ability to reconstruct high-resolution contact matrices from Pore-C and Micro-C data effectively. Applying these deep restoration models to Pore-C data in realistic settings represents a novel advancement that not only mitigates the inherent challenges posed by its complex and noisy nature but could significantly reduce sequencing costs for Pore-C experiments.

## Methods

Pore-C and Micro-C contact frequency matrices are produced from Pore/Micro-C experiments where thousands to millions of cells are processed at the same time. The result is a quadratic matrix *H*_*n×n*_ describing the contact frequencies between genomic positions at a given genomic bin *n*, which represents a certain genomic interval e.g. 1000kbp or 5000kbp. A higher the value *H*(*i, j*) indicates higher spatial proximity of *i, j* in the three-dimensional space of the nucleus.

### Pore-C preprocessing

The training dataset for Pore-C was collected from Deshpande et al., comprising HG002 genome NlaIII-digested Pore-C libraries sequenced using PromethION, and accessible from the Gene Expression Omnibus (GEO) under accession number GSM4490691 [21]. Raw fastq files were obtained from the Sequence Read Archive (SRA) database (Project: PRJNA627432, Experiment Accession: SRX8156771).

The fastq files were processed using the wf-Pore-C version 1.0.0 workflow from Oxford Nanopore, provided by epi2me labs (https://github.com/epi2me-labs/wf-pore-c). This workflow was employed to derive high-order chromatin contacts from Pore-C concatemers. It involved pre-processing the reference genome and fastq files, aligning reads to the GENCODE reference genome (GRCh38.p14 primary assembly), and post-processing the aligned reads (including filtering spurious alignments, detecting ligation junctions, and assigning fragments). The wf-Pore-C workflow generated final pairs files as output, serving as required inputs for the CCUT.

### Evaluation metrics

For the problems of image reconstruction and single image super-resolution, the loss functions and evaluation metrics aim to measure the distance between the predicted output of the model and the ground truth (high-resolution CC map). Metric calculation was applied after data augmentation. Since appropriate metrics are part of an active debate in the community, we will report the most common ones. Given a noise-free m×n monochrome image H and its noisy approximation K:

#### Mean Absolute Error / L1 Loss

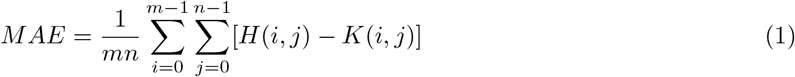

#### Mean Squared Error / L2 Loss

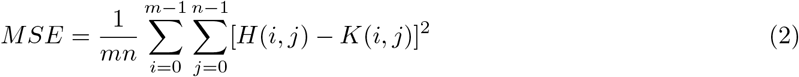

#### Huber Loss

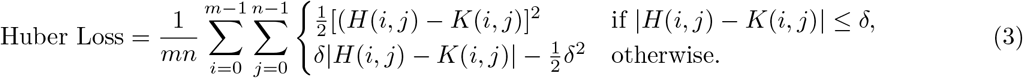

Where:

- *H*(*i, j*) represents the predicted value.
- *K*(*i, j*) represents the ground truth value.
- *δ* is a threshold parameter.

In this case, *δ* = 1.

#### Peak Signal-to-Noise Ratio

PSNR is defined via the MSE:

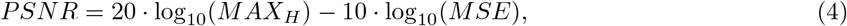

where *MAX*_*H*_ is the maximum pixel value of the image.

#### Structural Similarity Index

The Structural Similarity Index (SSIM) as a loss function is given by:

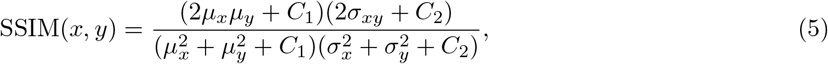

where

*μ*_*x*_ = mean of *x*,

*μ*_*y*_ = mean of *y*,

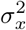 = variance of *x*,

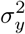 = variance of *y*,

*σ*_*xy*_ = covariance of *x* and *y*,

*C*_1_ = (*K*_1_*L*)^2^, and

*C*_2_ = (*K*_2_*L*)^2^ are constants.

*L* is the dynamic range of the pixel values, *K*_1_ = 0.01 and *K*_2_ = 0.03 by default.

The SSIM-based loss is then calculated as:

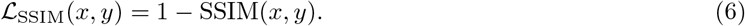

#### Fourier Transformed Losses

The toolbox also provides variations of the above losses combined with a loss incorporating the fourier transformations of the low-resolution and high-resolution images and is provided as an example based on the L1 and SSIM losses.

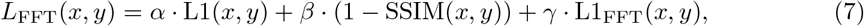

where

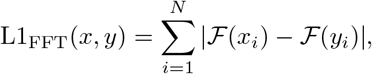

*α* = 0.1, *β* = 1.0, *γ* = 0.1 are the weights for loss components,

*N* is the number of pixels in the matrices,

*ℱ* denotes the Fourier transform.

#### Perceptual Loss

We integrated the Perceptual and VGG loss as implemented by ESRGAN into our framework [44]. Perceptual loss, often used in advanced image processing tasks, evaluates the perceptual difference between the target and the output images, rather than focusing purely on pixel-wise errors. This loss function leverages deep neural networks, typically pre-trained on image recognition tasks (ee utilize VGG [45]), to extract feature representations from intermediate layers. By comparing these feature representations of the predicted and ground truth images, Perceptual loss captures more nuanced discrepancies in texture, color, and structural details that are often overlooked by traditional loss functions.

#### Benchmark Model

We employed a U-Net-based architecture, integrated with Residual-in-Residual Dense Blocks (RRDB) derived from the Enhanced Super-Resolution Generative Adversarial Network (ESRGAN) framework [44], which is particularly adept at handling complex image reconstruction tasks. The model is configured with an input and output of 3 channels each, typically accommodating standard RGB images. It operates through a sequence of feature layers set by default to [64, 128, 256, 512, 1024], enabling a gradual compression and subsequent expansion of the input data. The downsampling pathway consists of a series of RRDB and convolutional blocks. Each RRDB layer is immediately followed by a convolutional block that not only adjusts the channel dimensions to the subsequent feature layer but also prepares the tensor for downsampling via a max pooling operation with a kernel size of 2 and a stride of 2, followed by a bottleneck layer doubling the number of features. For upsampling, the model employs transposed convolutions for each feature level, effectively doubling the spatial dimensions of the tensor. Post-upsampling, RRDB blocks are applied, which enhances the feature refinement without altering the tensor size. Each upsampling step also includes a concatenation with the corresponding skip connection from the downsampling path, followed by another convolutional block to merge the features seamlessly.

### Model Training

All models discussed in this study were trained using an NVIDIA RTX 3090 GPU with xx GB vRAM. For the main benchmark with unprocessed 3C data, each model underwent training for 10 epochs utilizing the Adam optimizer, which was selected for its effectiveness in various machine-learning contexts. The optimizer was configured with a learning rate of 0.0005, a setting determined to balance convergence speed and stability. Additionally, to maintain the integrity of the model updates and to optimize computational efficiency, gradient clipping was implemented alongside the strategic use of half-precision computations where applicable. Early stopping was applied when models did not improve in MSE over five epochs. All models were trained on the gold standard Micro-C dataset from Krietenstein et al.(4DN: 4DNES2M5JIGV, [5]). This dataset was specifically processed at a resolution of 10k bp. For the Pore-C reconstructions, we used the gold standard dataset from Deshpande et al (GEO: GSM4490691, [46]. The Pore-C dataset was processed at a higher resolution of 50k bp. Each training dataset comprised slices of 200×200 pixels, encapsulating data from chromosomes 1 through 18. This selection was made to ensure a comprehensive representation of the genome, while chromosomes 19 to 22 were reserved exclusively for model evaluation purposes, enabling a separate and unbiased assessment of model generalizability and performance.

## Data Availability

For this study, we used a Pore-C and Micro-C dataset. The gold standard Micro-C dataset by Krietenstein et al. was obtained in pairs format from 4DN: 4DNES2M5JIGV [5]. The Pore-C data from Deshpande et al was obtained from GEO: GSM4490691 and processed with the Oxford Nanopore best practices Nextflow pipeline: wf-pore-c to pairs format. We downsampled the data by their respective factors in the pairs format since it is lossless and contains read-level information and afterward constructed the multi-resolution contact maps using the Cooler library and following their best practice guideline for pairs data[47].

## 1 Code Availability

All scripts and the tool can be accessed via GitHub.

## Acknowledgements

We acknowledge funding from the Mainz Institute of Multiscale Modeling and from the Emergent AI Center funded by the Carl-Zeiss-Stiftung. SG, SS, AK and ACN acknowledge funding by SFB 1551 Project No. 464588647 of the Deutsche Forschungsgemeinschaft (DFG). KES acknowledges funding from the German Research Foundation (DFG) Project No. 320163632. We thank Kyra Klos and Robin Msiska for the discussions.

